# Bidirectional FtsZ filament treadmilling transforms lipid membranes via torsional stress

**DOI:** 10.1101/587790

**Authors:** Diego A. Ramirez-Diaz, Adrian Merino-Salomon, Fabian Meyer, Michael Heymann, German Rivas, Marc Bramkamp, Petra Schwille

## Abstract

FtsZ is a key component in bacterial cell division, being the primary protein of the presumably contractile Z ring. *In vivo* and *in vitro*, it shows two distinctive features that could so far however not be mechanistically linked: self-organization into directionally treadmilling vortices on solid supported membranes, and shape deformation of flexible liposomes. In cells, circumferential treadmilling of FtsZ was shown to recruit septum-building enzymes, but an active force production remains elusive. To gain mechanistic understanding of FtsZ dependent membrane deformations and constriction, we designed an in vitro assay based on soft lipid tubes pulled from FtsZ decorated giant lipid vesicles (GUVs) by optical tweezers. FtsZ actively transformed these tubes into spring-like structures, where GTPase activity promoted spring compression. Operating the optical tweezers in lateral vibration mode and assigning spring constants to FtsZ coated tubes, we found that FtsZ rings indeed exerts 0.14 – 1.09 pN forces upon GTP hydrolysis, through torsional stress induced by bidirectional treadmilling. These directional forces could further be demonstrated to induce membrane budding with constricting necks on both, giant vesicles and *E.coli* cells devoid of their cell walls.

## Main Text

In biology, fundamental mechanical processes, such as cell division, require an intricate space-time coordination of respective functional elements. However, how these elements, mostly proteins, can self-organize to exert forces driving large-scale transformations is poorly understood. In several organisms, ring-like cytoskeletal elements appear upon cytokinesis; for instance, the FtsZ-based contractile Z ring in bacteria. Ring-like FtsZ structures have previously been shown to deform liposome membranes (1,2). When reconstituted on flat membranes, FtsZ self-assembles into rotating-treadmilling vortices with conserved direction (3,4). *In vivo*, FtsZ shows circumferential but bidirectional treadmilling that is assumed to serve as a pacemaker guiding peptidoglycan synthesis around the septum (5,6).

Despite of these exciting findings, it is not clear whether these treadmilling FtsZ filaments actively contribute to the physical process of lipid membrane constriction and cytokinesis in bacteria (7,8). The challenge is two-fold: (i) how much force is actually required to divide bacteria, given the mechanical coupling of the lipid membrane and the cell wall?, and (ii) even if FtsZ filaments can generate membrane deforming forces, what is the exact mechanism by which these forces are exerted? For instance, considering the mechanical bearing related to internal turgor pressure (~MPa), models have suggested that FtsZ forces in the range of 8-80 pN would be required for constriction (9). In contrast, it has been proposed that turgor pressure need not be considered, due to the possibility of same osmolarity between periplasm and cytoplasm (10). For this case, very low FtsZ forces in the range of 0.35 – 2.45 pN could exert membrane deformations leading to constriction (10). In conclusion, *in vivo* and *in vitro* experimental approaches addressing those two major questions are needed to gain deeper understanding in cell division in bacteria.

Here, we have employed *in vitro* reconstitution as a strategy to understand the mechanistic features of FtsZ as a membrane deforming polymer. Using an optical tweezers-based approach by pulling soft lipid tubes from deflated giant unilamellar vesicles GUVs, our aim is to quantitatively elucidate the physical principles underlying membrane deformations induced by dynamic FtsZ rings on GUVs and the scale of delivered forces. These particular principles are key to understand the nature of FtsZ membrane deformations *in vitro* and *in vivo*.

Based on our recent study (3), we externally added FtsZ-YFP-mts to GUVs made of *E. coli* lipid extract. Conditions to obtain ring-like structures were determined by tuning GTP and Mg^+2^ (Fig. 1A). Since no clear deformations were observed for tensed vesicles (Fig. 1A), we designed a two-side open chamber allowing for slow water evaporation to obtain deflated and deformable GUVs. After 20-30 minutes, we evidenced that rings were inducing inwards-cone structures emerging from the membrane surface, indicative of drilling-like inward forces (Fig. 1B). Motivated by this specific geometry, we designed PDMS microstructures mimicking such inward cones (Fig. 1C Fig 1SA). After coating these with supported lipid bilayer (SLB) and triggering protein polymerization, we observed individual filaments/bundles to wrap the cone in a dynamic fashion resembling a vortex (Fig. 1D) (Movie S1). We noticed that the dynamic vortices rotate both clockwise and anticlockwise (Fig. 1E), indicating that preferential directionality observed on flat SLBs is absent in conical geometry. Rotational velocities were estimated around 43 *nm*/*s*, showing relatively good agreement with our previous results on flat surfaces (34 *nm*/*s*) (3).

**Figure 1.**
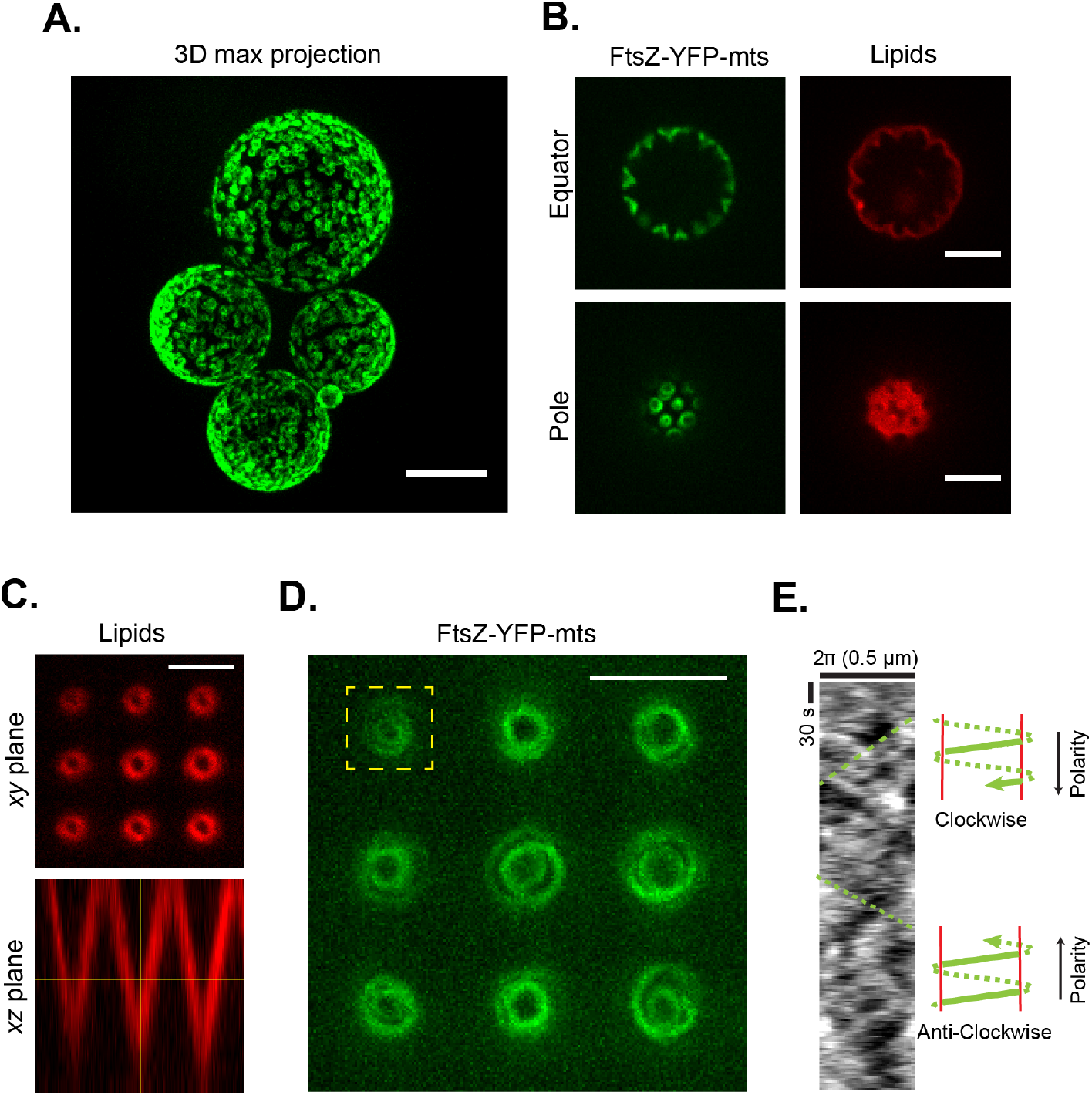
**A)** FtsZ-YFP-mts ring structures externally decorating GUVs (scale bar=10 μm). **B)** After GUV deflation, inwards conical deformations emerged from FtsZ rings. **C)** Inspired by deformations in (B), we designed a PDMS microstructure with inwards-conical geometry covered with a supported lipid bilayer (SLB). The imaging plane was chosen to have a cross-section of ~1 μm diameter. **D)** Inside cones, FtsZ-YFP-mts self-assembled into dynamic vortices (Movie S1). **E)** Kymograph showed negative and positive slopes indicating the presence of clockwise and anticlockwise directions (Scale bar = 5 μm).

To quantitatively characterize the impact of FtsZ on soft tubular geometries, we developed a method based on optical tweezers. Contrary to prior approaches using micropipettes (11), we pulled soft tubules from weakly surface-attached GUVs (Fig. 2SA) by moving the GUVs relative to an optically trapped bead. Lipid tubes with mean diameter of ca. 0.47 *μm* (Fig. 2SB) were now pulled from deflated GUVs decorated with ring-like FtsZ structures and inward-conical deformations (Movie S2). Once tubes were formed, protein filaments entered and deformed the tube. After 175s, helical tube shapes were clearly observed (Fig. 2A), indicative of dynamic coiling (Movie S3). As more protein entered the tube and accumulated in the tip, the spring-like structure got compressed (Fig. 2A, 500s). These helical tube deformations can be rationalized by twisting of an elastic rod subjected to constant tensile force (Fig. 3F). Similar to the experiment in Figure 1D, filaments grew towards (clockwise) and away from (counterclockwise) the tip of the tube. If filament growth imposes torsion, the counter-growing filament will generate torsion in the opposite direction. These two different torsional contributions result in the buckling of the lipid tube and the formation of a 3D helix (Fig. 3F). The importance of the bidirectional treadmilling, or bidirectional filament growth, can be understood using a shoelace analogy: opposite torque should be exerted on both ends of the shoelace to observe a helical deformation. If one end is loose, the opposite end will only rotate accordingly (sliding). A net force due to FtsZ twisting and coiling in the outwards-vesicle direction caused the incorporation of new lipid material from the flaccid vesicle to the tube.

**Figure 2.**
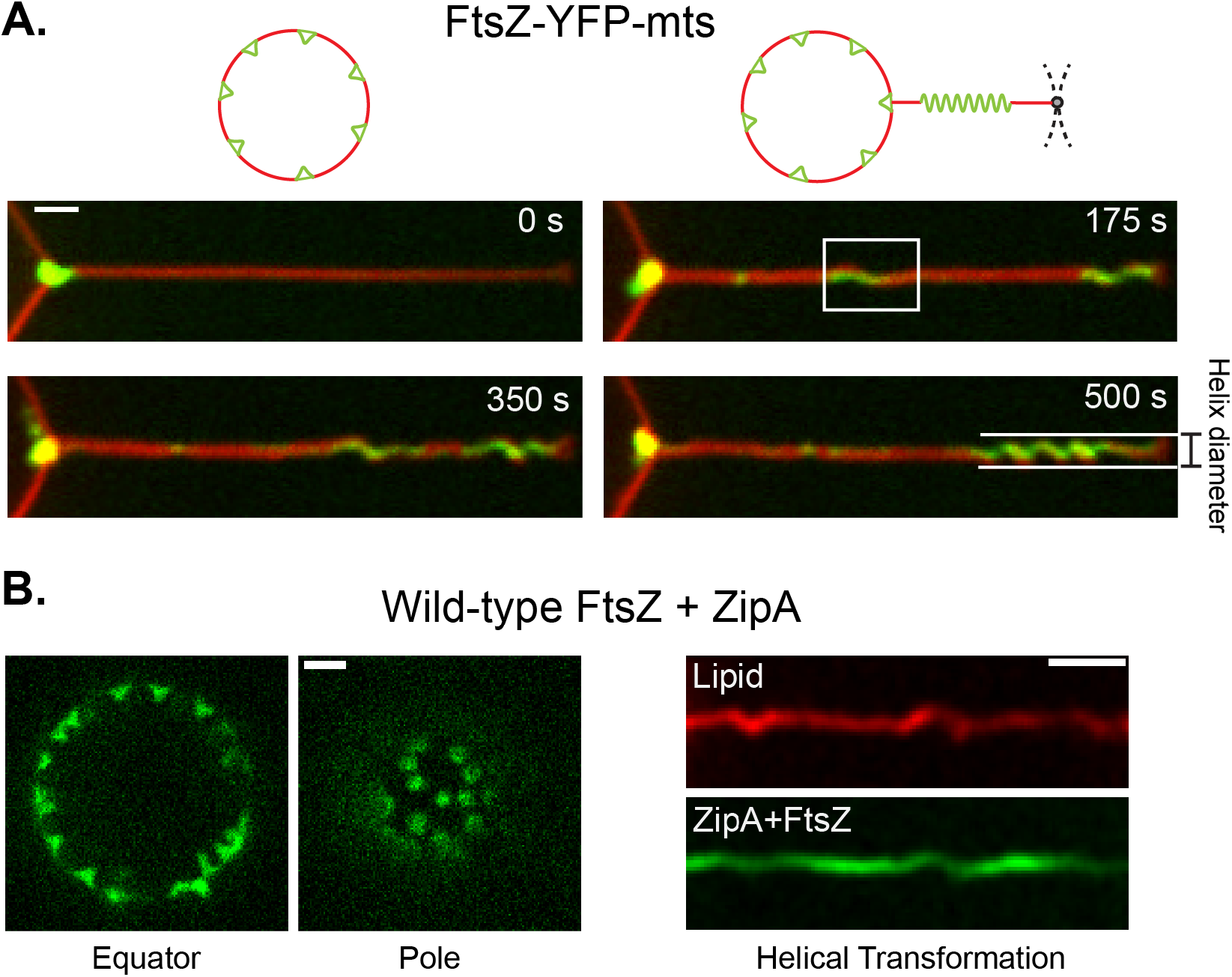
**A)** To understand inwards deformations, we stretched the cone structures into a tubular geometry. Soft lipid tubes were pulled from weakly surface-attached GUVs by moving the GUVs relative to an optically trapped bead. As long as the FtsZ-YFP-mts entered the tube, a process of coiling is clearly observed as a function of time. As a result, protein being accumulated over the tip transformed the lipid tube into a spring-like shape. B) To rule out that artificial attachment of the FtsZ-YFP-mts is responsible of the helical transformation, we reconstituted wild-type FtsZ anchored to the membrane via ZipA to obtain ring-like structures decorating GUVs. When vesicles were deflated, wild-type FtsZ + ZipA caused inwards cone-like deformations. After pulling lipid tubes, similar helical deformations were observed confirming that torsion is related the FtsZ core of the polymer. Fluorescence signal of wt-FtsZ-Alexa 488 is shown in green while siZipA remains unlabeled. (Scale bar = 2 μm).

**Figure 3.**
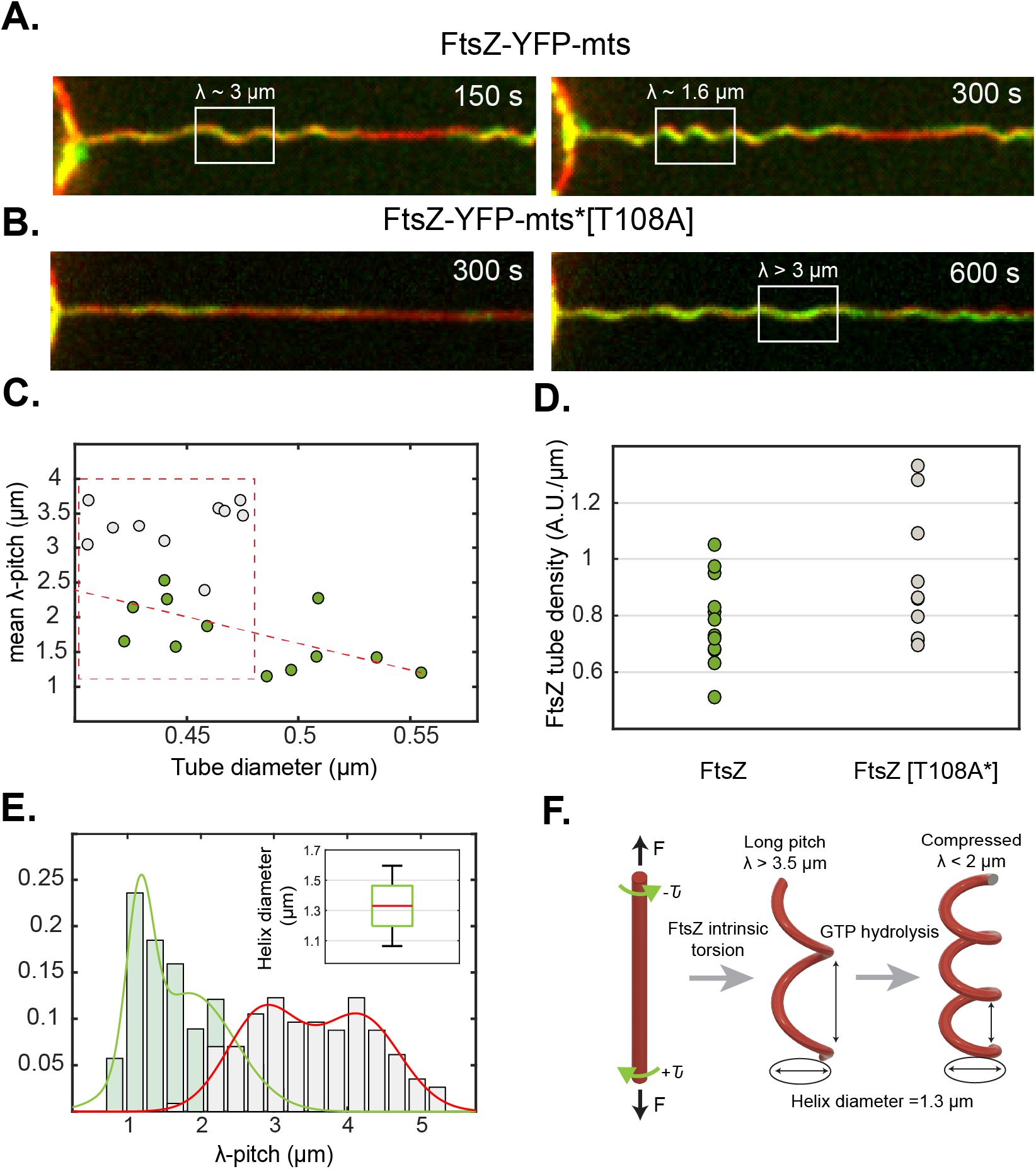
**A)** FtsZ-YFP-mts and **B)** FtsZ-YFP-mts*[T108A] promoted helical deformations with the difference that GTPase activity induce compression (λ ~ 1.6 μm) of initially longer pitch (λ > 3 μm). **C)** To rule out that compression was biased by the deflation state, we plotted tube diameter vs mean pitch for FtsZ-YFP-mts (N=12) (green) and FtsZ-YFP-mts*[T108A] (N=10) (gray). Despite of higher tube densities for FtsZ-YFP-mts*[T108A] as shown in **(D),** the mean pitch for no GTPase case is longer at comparable tube diameters. **E)** We observed two clear pitch states for FtsZ-YFP-mts (gray bars/green line) and FtsZ-YFP-mts*[T108A] (gray bars/red line) with a clear dominance of longer pitch for the mutant without GTPase activity. **F)** Helical deformations can be understood by twisting an elastic rod subjected to constant force. We postulate that FtsZ has an intrinsic torsion that is enhanced by GTPase activity, driving further compression. Intrinsic FtsZ torsion rules long-pitch transformations (λ > 3 μm) while GTP enhances further torsion causing higher pitch states (λ < 2 μm).

Since spring-like deformations were observed with a FtsZ protein chimera that binds autonomously to membrane (FtsZ-YFP-mts), we attempted to confirm whether this phenomenology is intrinsic to the FtsZ polymer and not caused by the membrane targeting sequence. Based on the reconstitution of dynamic rings on flat membranes using the *E.coli* FtsZ natural anchor ZipA (12), we identified the right conditions to obtain WT-FtsZ rings externally decorating GUVs using ZipA (Fig. 2B). In the same way as for FtsZ-YFP-mts, rings induced inwards cone-like deformations on deflated vesicles (Fig. 2B). The obvious next step was to evaluate their impact on a soft tubular geometry. As a control, we pulled lipid tubes having only ZipA to evidence missing deformation. Then we added WT-FtsZ and observed helical transformations as expected (Fig. 2B), indicating that FtsZ polymer and not its membrane attachment caused this effect. Interestingly, FtsZ-YFP-mts as well as FtsZ+ZipA displayed in plectonic/supercoiled regions (Fig. 1SE) as further indicative of torsion over the lipid tube.

To investigate the role of GTP hydrolysis, we reconstituted FtsZ-YFP-mts*[T108A], a mutant with low GTPase activity (3). We observed that FtsZ-YFP-mts*[T108A] also self-assembled into ring-like structures (Fig. 1SB) that lacked dynamic treadmilling (3) yet still promoted inwards deformations (Fig. 1SC). Interestingly, the activity of FtsZ-YFP-mts*[T108A] on the tubes was much delayed (Fig. 2SC). Although helical deformations were also observed after 350s (Fig. 3B), their pitch remained considerably longer (*λ* > 3 *μm*) also at long times (900s). In contrast, helices decorated with GTP-active FtsZ-YFP-mts (Fig. 3A) underwent compression to a pitch of *λ* ∼ 1.5 *μm* after 300s. By plotting the arc-length of the spring against FtsZ density on the tube, we clearly observed a greater membrane-deforming activity for FtsZ-YFP-mts (Fig. 2SE). Experiments shown in Fig. 3A-B correspond to similar tube diameters (d = 0.44 μm Fig. 2SB).

Since the deflation of individual GUVs could vary, we also tested whether compression could be biased by GUV membrane tension (deflation) and protein density over the tube. The tube diameter *d* represented our observable for membrane tension according to the relation 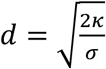, where *κ* denotes the lipid bending modulus and *σ* the membrane tension (11,13). The lower the membrane tension (deflation), the larger the tube diameter. Therefore, we plotted the mean pitch vs tube diameter (Fig. 3C), for independent experiments, considering also the amount of protein (Fig. 3D). Although there was a mild correlation between pitch and diameter (Fig. 3C) for FtsZ-YFP-mts, the mean pitch was consistently longer for FtsZ-YFP-mts*[T108A] (Fig. 3C) in the case of tubes with comparable or higher protein density (Fig. 3D). To better visualize the impact of GTPase activity, we plotted the pitch distribution for both proteins: the GTPase activity contributed to a decrease of pitch (Fig. 3E) as clear indicative of spring compression. Interestingly, both distributions are reasonably bimodal, indicative of two states of torsion: a structural intrinsic torsion (longer pitch) that is further enhanced (shorter pitch) via GTPase activity (Fig. 3F). Note that FtsZ-YFP-mts*[T108A] could exhibit residual GTPase activity driving some compression.

To quantitatively characterize the mechanical properties of FtsZ-YFP-mts-induced spring-like structures, we implemented an alternative approach based on the elastic response of the GUV + tube to a specific dynamic input. Using a piezoelectric stage, we induced a lateral oscillation of the GUV position (*A* = 3 *μm*, *f* = 1 *Hz*) and recorded forces by the optical trap (Movie S4). We here measured the resistive force of the material per micrometer (k-spring constant). The stiffer the material, the higher force detected by the optical trap. To calculate the amplitude of the signal at 1 Hz the signal was Fast Fourier Transformed, as depicted in Figure 4B, where the red line refers to the pure lipid tube and the green line to lipid + FtsZ. Due to variability in terms of vesicle size, deflation state and FtsZ surface concentration, a range of values was here reported rather than a normal-distribution. The pure lipid contribution (N=11) had values between 0.15 – 0.55 pN/ *μm* (Fig. 4C), while for the lipid + FtsZ system (Fig. 4A) (N=36) we determined values between 0.23 – 1.52 pN/ *μm* (Fig. 4C). Note that for some vesicles, the pure lipid response yet dominated the spring constant measurement. Based on these results, we next attempted to estimate the range of forces that a single FtsZ ring delivered. To this end, the FtsZ fluorescence signal per single ring was determined and compared to the signal on the FtsZ-covered tubes (Fig. 2SF). Thus, for each force measurement we were able to approximate the “number of rings” according to the total FtsZ brightness. By plotting these forces vs “number of rings” (Fig. 4D), we noted that scattered data can be well described by two straight lines, and consequently their slopes defined the range of forces per ring: 0.14-1.09 pN.

**Figure 4.**
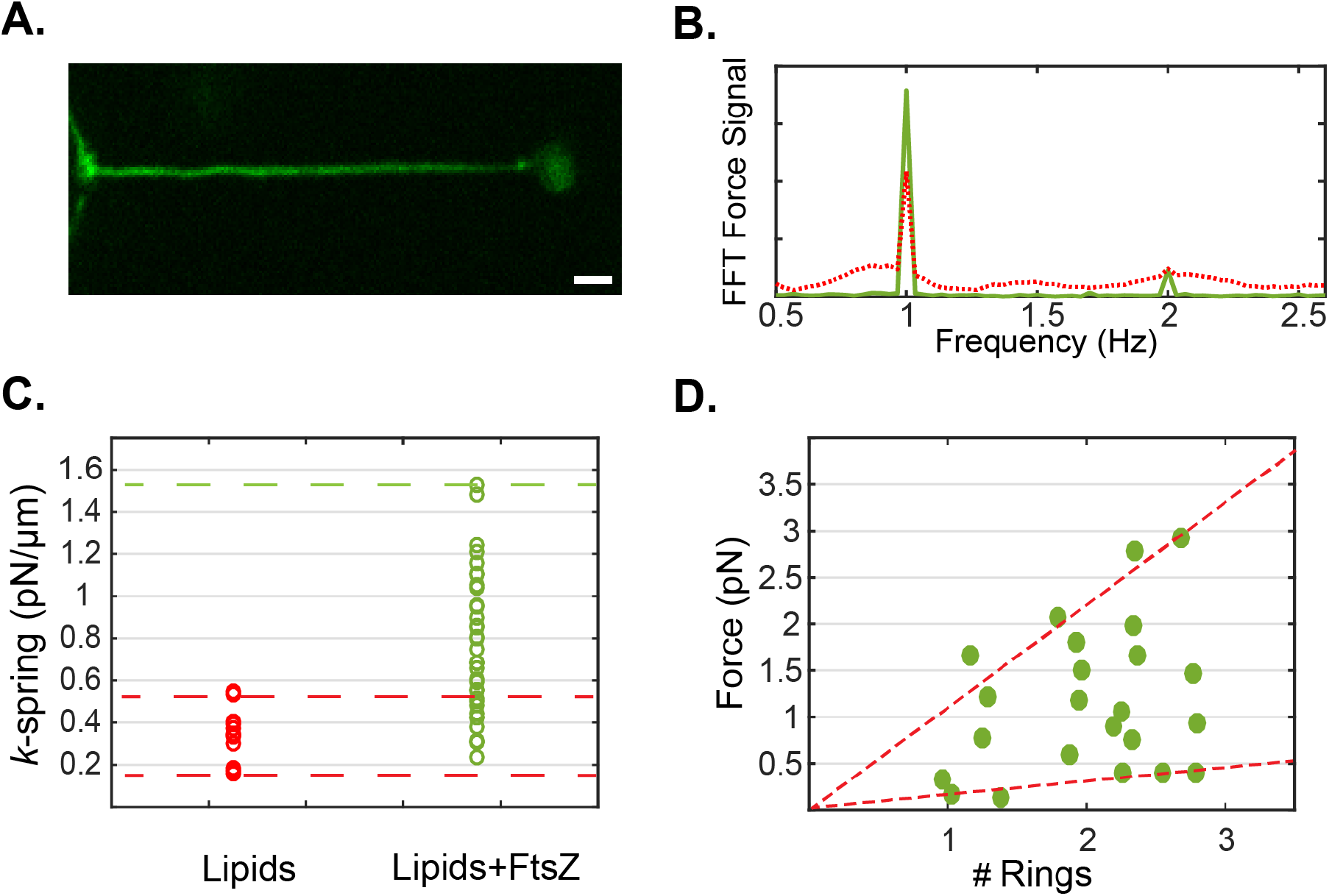
**A)** Spring-like structures were mechanically assessed by forcing the tube length to oscillate with an amplitude of 3 μ*m* and a frequency of 1*Hz*. **B)** To measure forces, we tracked bead-displacement as response of the dynamic input. Then, we calculated the Fast Fourier Transform (FFT) to calculate the amplitude of the signal. Red line: lipid signal and green line: FtsZ. **C)** By calculating the amplitude, we assessed the spring constant for the case of the only lipid contribution (N=11) and Lipid+FtsZ (N=36). Dashed red lines indicate the range where the lipid response dominated over the FtsZ contribution to the spring constant. **D)** By assessing the average total intensity of FtsZ rings on a flat surface with the same imaging conditions, we estimated the number of rings for each FtsZ experiments shown in **C)**. Therefore, scattered data representing force vs. number of rings could be described by two dashed red lines with slopes 0.14 and 1.09 pN per ring. These slopes represented the upper and lower limit for the FtsZ forces per ring.

Interestingly, we had previously inferred that FtsZ-YFP-mts rings on SLB are made of filaments of ~0.39 *μm* in average (3). This estimation could be used to validate our force measurements. A FtsZ filament with a persistence length ~0.39 *μm* exhibits a flexural rigidity *K* = 1.59 × 10^−27^ *Nm*^2^ that agrees well with Turner et. al., (2012) (14). Based on this, we could calculate the Young’s modulus of FtsZ filaments: *E*_*FtsZ*_ = 51.8 *MPa*. (*E* = *KI* where *I* = *πr*^4^/4, the area moment of inertia, *r* = 2.5 nm (15)). On the other hand, the Young’s modulus *E* of a spring is related to the spring constant through *E* = (*k l*_0_)/*S*, where *k* denotes the spring constant, *l*_0_ is the spring initial length and *S* the cross-section. Since *l*_0_/*S* was fairly constant in our tube experiments, we could claim that *E*_*FtsZ*_/*E*_*l*_ = (*k*_*FtsZ*_/*k*_*l*_) . To calculate 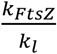, we here considered raw averages for distributions shown in Fig. 4C and subtracted the lipid contribution in the case of FtsZ: *k*_*l*_ = 0.34 pN/ *μm* and *k*_*FtsZ*_ = 0.6 pN/ *μm*. Then, the ratio 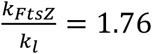 showed good agreement compared to 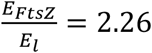 assuming *E*_*l*_ = 22.9 MPa (lipids with a bending *κ* = 20 *k*_*b*_*T*) (16, 17). This confirmed that our force measurements corresponded well with previous flexural rigidity values for FtsZ fibers. In addition, our data provide further evidence that FtsZ filaments are softer than other cytoskeleton proteins such as microtubules (*K*~10^−23^) or actin (*K*~10^−26^) (18,19)

The helical nature of FtsZ and its torsional dynamics have been experimentally observed (20, 21); however, its relation to a potential mechanism of deforming membranes has not yet been clearly established. According to our observations, the helical membrane transformation caused in this study by FtsZ filaments can best be understood by assuming Darboux torque around the lipid tube. Darboux torques are tangential torques caused by a local mismatch between the plane defined by the filament curvature and the membrane attachment direction (22). This twisting angle along the one filament is key to produce torque. A molecular dynamics study showed that dynamin, a helical endocytic constriction protein, required twisting of the “adhesive-stripe" to achieve full membrane hemifusion (22). In the case of FtsZ, molecular dynamics studies have predicted an angle of “twisting” along the c-terminus, where membrane attachment occurs (23, 24). Also, Fierling and coworkers have theoretically studied membrane deformations produced by filaments inducing torques (25). Strikingly, they found inward vortex-like deformations from flat surfaces and spring-like shapes when filaments wrapped around a tubular geometry (25). These predictions agree remarkably well with our observations.

So far, we had investigated an inverse geometry, i.e., FtsZ added from the outside, as compared to the physiological case. Now we also reconstituted FtsZ-YFP-mts and FtsZ-YFP-mts*[T108A] inside GUVs (Fig. 5A). Conditions to obtain ring-like-structures (Fig. 5B) or filaments wrapping the vesicle (Fig. 1SD) were again found by tuning GTP and Mg^+2^. Interestingly, the diameters of FtsZ-YFP-mts*[T108A] rings were significantly larger (0.89 *μm*) than FtsZ-YFP-mts (0.44 *μm*) (Fig. 5D). This difference was not observed in the case of SLBs (3), indicating the possibility that softer lipid surface affects the steady state of FtsZ assembly. In other words, the physical properties of the lipid surface, such as stiffness, may play an important role in FtsZ fragmentation and treadmilling. In addition, the wide size distribution in the low GTPase mutant case (Fig. 5D) implied that polymers were flexible to accommodate a larger variety of curvatures. Strikingly, both FtsZ mutants could create outwards deformations emerging from rings (Fig. 4E). But only in the case of FtsZ-YFP-mts, there was clear evidence of constricting rings (Fig. 4E) similar to previous reports (1). Based on Fig. 1, we hypothesize that FtsZ torsion could create outwards out-of-plane forces (Fig. 5F). However, FtsZ filaments only exhibiting static (structural) torsion were unable to stabilize smaller diameters. In contrast, dynamic twisting upon GTP-hydrolysis drives constriction and clustering (Fig. 5G), such that active FtsZ filaments lead to an overall shrinkage of diameters. FtsZ constriction and neck formation on flat membranes thus represents an analog of helix compression, where forces of 0.14 – 1.09 pN per ring are actuating the lipid surface.

**Figure 5.**
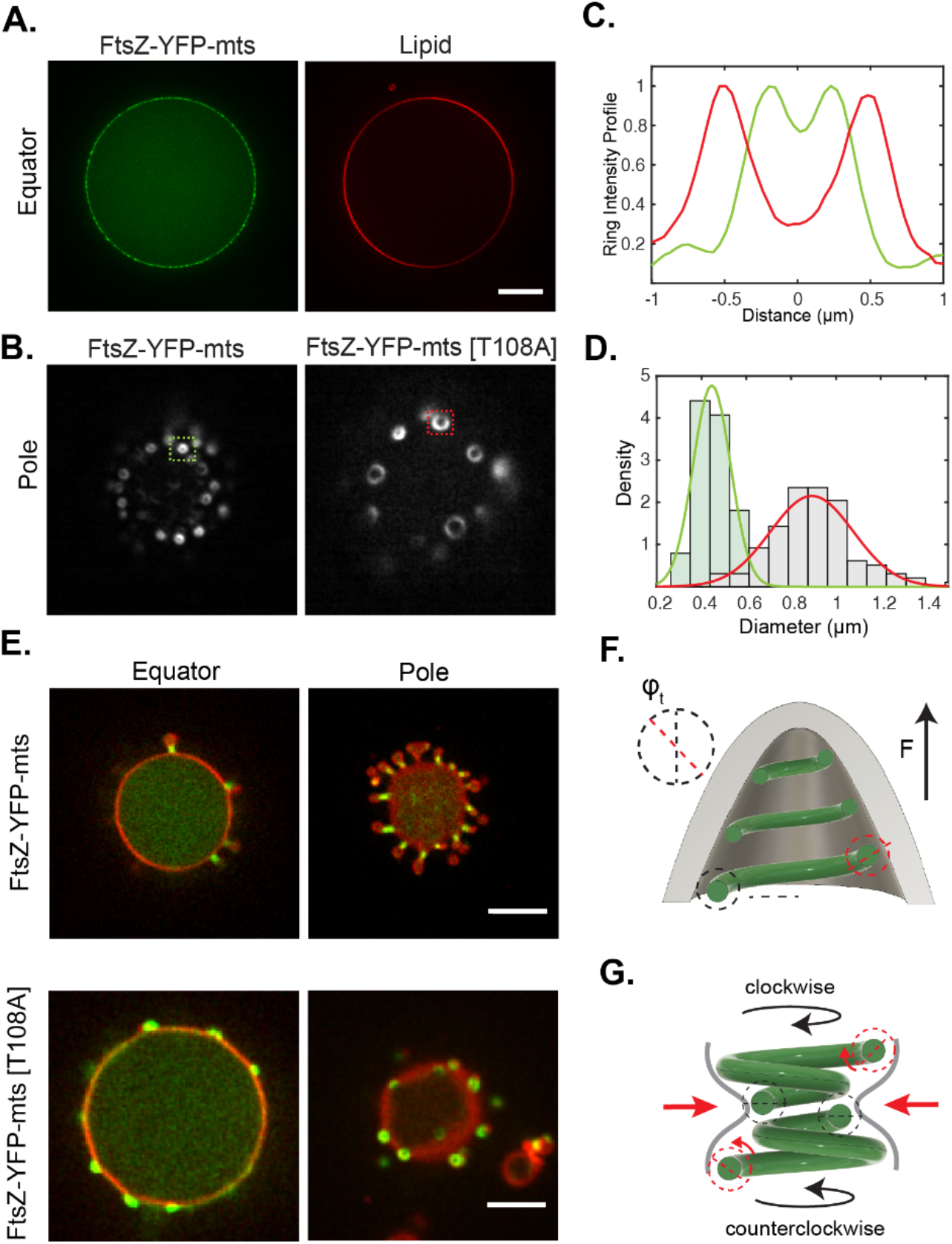
**A)** FtsZ-YFP-mts and FtsZ-YFP-mts*[T108A] rings inside GUVs. **B)** Imaging of rings, at GUVs bottom, using TIRF microscopy. **C)** Intensity profile of structures indicated in (B) showed that FtsZ-YFP-mts (green line) rings exhibit smaller diameter than FtsZ-YFP-mts*[T108A] (red line). **D)** Size distribution of (N=112) FtsZ-YFP-mts*[T108A] (gray bars and red line) and (N=102) FtsZ-YFP-mts showed a drastic reduction in ring diameter due to GTP hydrolysis. **E)** After deflation, both mutants drove outwards deformations with the difference that GTPase activity promotes constriction and neck formation. **F)** We suggest that intrinsic torsion can create out-of-plane forces; however, **G)** GTP hydrolysis triggered a super-constricted state favoring higher curvatures.

In order to appreciate the relevance of our mechanistic findings for lipid membranes *in vivo*, we needed to transfer our FtsZ-YFP-mts construct to the cellular environment. Therefore, *E. coli* cells were cloned and transformed with the corresponding gene into an inducible plasmid. Upon IPTG induction, FtsZ-YFP-mts fluorescence signals in the cells were observed. The FtsZ-YFP-mts construct localizes in several ring-like structures around midcell (Fig. 6A). Multiple Z-ring structures were observed, due to the overexpression of the FtsZ-YFP-mts protein (1). A 3D-reconstruction reveals that these FtsZ assemblies are indeed ring structures that resemble those formed by native FtsZ rings at the division site (Fig. 6A). Importantly, without addition of inductor, no FtsZ-YFP-mts structures were observed (Fig. 6B). Since FtsZ driven deformations were not observed for tensed GUVs (Fig. 5), we reasoned that in walled bacteria with turgor pressure, it might be difficult to observe FtsZ-YFP-mts driven membrane deformations. Therefore, cells were treated with lysozyme to create *E. coli* spheroplasts in osmoprotective medium. Cells expressing the FtsZ fusion protein were highly fragile and prone to lysis. We therefore started microscopic analyses before all cells have converted to spheroplasts (Fig. 6B). Importantly, vesicular structures budding out from spheroplasted cells were observed (Fig. 6B, arrows). These vesicular structures were not observed in control cells lacking the FtsZ-YFP-mts expression, indicating that they are a consequence of protein overproduction. We also observed drastic deformations of the plasma membrane that resemble plasmolysis. In these cases, FtsZ-YFP-mts assemblies underneath the membrane seem to pull in the membrane and exert force leading to a separation of plasma membrane and outer membrane. A membrane stain reveals that areas with strong FtsZ fusion protein assemblies also show membrane invaginations or constriction necks (Fig. 6C, arrows; Movie S5). These results agreed remarkably well with our outwards deformations and constriction necks from FtsZ rings inside GUVs. Directional screw-like forces promoting extrusion of lipid material or budding (Fig. 5H & Fig. 6B), as well as constriction necks (Fig. 5G & Fig. 6C), are both explained in terms of a FtsZ polymer able to exert torsional stress as explained above. Interestingly, this opens the possibility of FtsZ filaments playing an active role in cell division organisms that divide by budding, such as Acholeplasma laidlawii (28).

**Figure 6.**
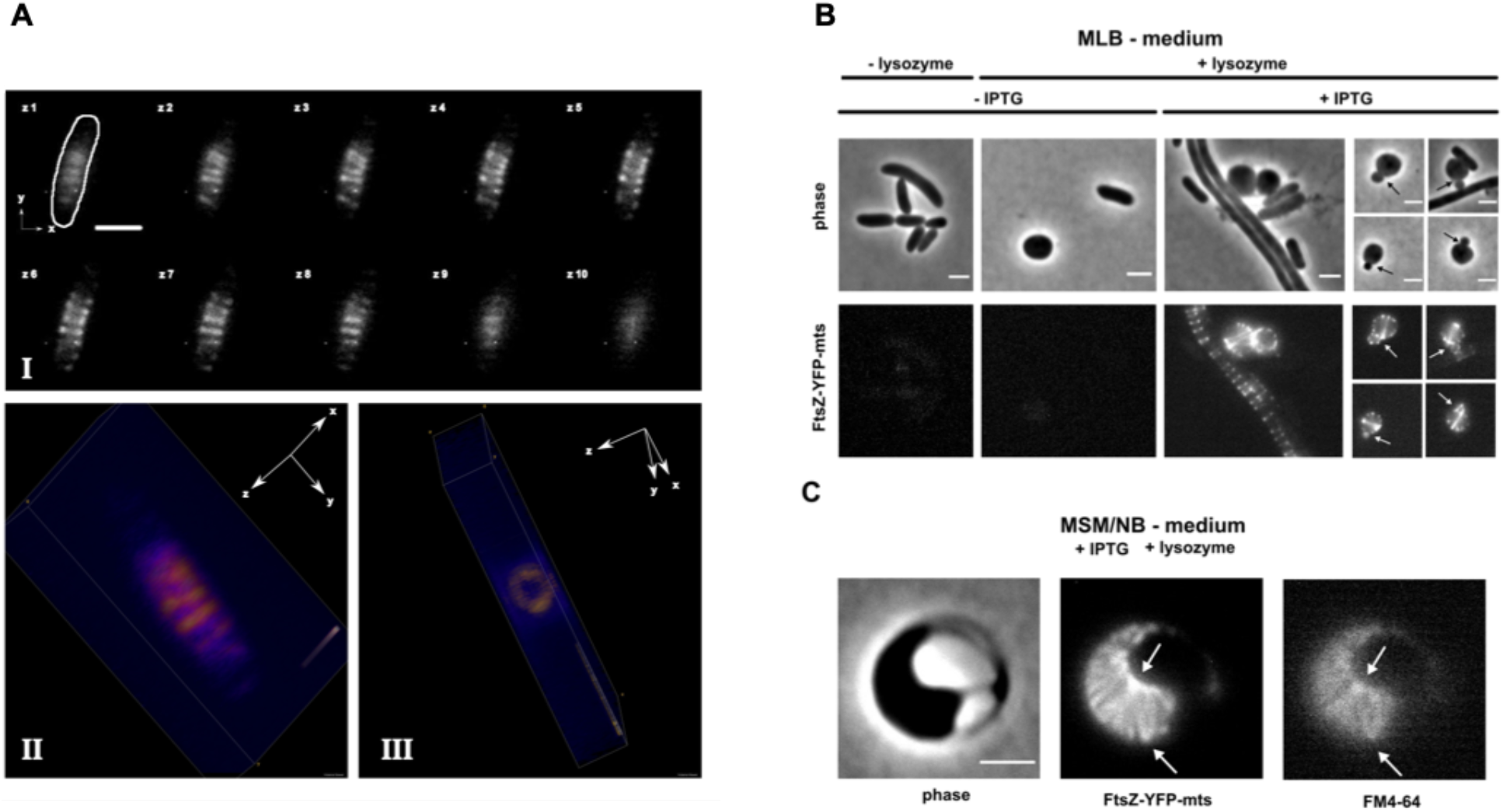
**A**) *E. coli* DH5α cells expressing FtsZ-YFP-mts polymeric structures perpendicular to the cell diameter around midcell (AI). 3D rendering reveals ring-like structures (AII-AIII). **B**) Removal of the cell wall by lysozyme treatment leads to spheroplast formation. Expression of FtsZ-YFP-mts leads to membrane vesiculation (arrows). **C**) Sphaeroplasts expressing FtsZ-YFP-mts show drastic deformations of the plasma membrane. FtsZ assemblies lead to local membrane invaginations (white arrows), indicating force generation. Scale bars 2 μm.

Altogether, our experiments provide clear evidence that FtsZ induces two kinds of mechanical deformations to membranes. Static (structural) FtsZ torsion rules the assembly of rings on flat surfaces (3) and induces inwards/outwards deformations as described earlier. Importantly, circumferential treadmilling powered by GTP hydrolysis induces an additional torque-twist, stabilizing smaller ring diameters and supporting further membrane constriction. Regardless whether protein was externally added or encapsulated, cylindrical geometry allowed clockwise and anti-clockwise treadmilling (Fig. 5G). Together active FtsZ induces helical transformation of the membrane tube and a super-constricted state of filaments, imposing a mechanical strain that promotes breakage and therefore the emergence of treadmilling (Fig. 5F). This establishes an interesting similarity between FtsZ and dynamin, in which GTP hydrolysis also triggers a super-constricted state, favoring fragmentation and clustering (19,20). We conclude that these torques represent a robust constriction mechanism for cylindrically shaped membranes, generating forces in the range of 0.14-1.09 pN per ring in the case of FtsZ. These FtsZ-induced forces drive outwards deformations and constriction necks in the case of deflated vesicles *in vitro* and wall-less *E. coli in vivo*. Although the here reported forces do likely not suffice for the entire process of bacterial cytokinesis of walled rod-like cells, given the temporal relevance of FtsZ dynamics in the coordination of synthesis of new wall material (5,6), an initial inwards membrane deformation may be key to trigger cytokinesis, in the form of a “curvature trigger”.

In view of our data, we hypothesize that if membrane tension is lowered; for instance, through the incorporation of de novo synthesized lipids in bacteria septum (10), the here reported force range might become relevant for the initiation of cell division.

## Acknowledgments

We acknowledge Dr. Sven Vogel and Dr. Allen Liu for very useful discussions. We thank MPIB Core Facility for assistance in FtsZ-YFP-mts protein purification. Funding through MaxSynBio consortium (Federal Ministry of Education and Research of Germany and the Max Planck Society) grant number 031A359A. The authors have declared that no competing interests exist. All data is available in the main text or the supplementary materials.

## Code availability

All custom code is available on request.

## Data availability

All data are available in the main text, the supplementary materials, or upon request.

